# A complex resistance locus in *Solanum americanum* recognizes a conserved *Phytophthora* effector

**DOI:** 10.1101/2020.05.15.095497

**Authors:** Kamil Witek, Xiao Lin, Hari S Karki, Florian Jupe, Agnieszka I Witek, Burkhard Steuernagel, Remco Stam, Cock van Oosterhout, Sebastian Fairhead, Jonathan M Cocker, Shivani Bhanvadia, William Barrett, Chih-Hang Wu, Hiroaki Adachi, Tianqiao Song, Sophien Kamoun, Vivianne GAA Vleeshouwers, Laurence Tomlinson, Brande BH Wulff, Jonathan DG Jones

## Abstract

Late blight caused by *Phytophthora infestans* greatly constrains potato production. Many *Resistance (R)* genes were cloned from wild *Solanum* species and/or introduced into potato cultivars by breeding. However, individual *R* genes have been overcome by *P. infestans* evolution; durable resistance remains elusive. We positionally cloned a new *R* gene, *Rpi-amr1*, from *Solanum americanum*, that encodes an NRC helper-dependent CC-NLR protein. *Rpi-amr1* confers resistance in potato to all 19 *P. infestans* isolates tested. Using association genomics and long-read RenSeq, we defined eight additional *Rpi-amr1* alleles from different *S. americanum* and related species. Despite only ∼90% identity between Rpi-amr1 proteins, all confer late blight resistance but differentially recognize *Avramr1* orthologs and paralogs. We propose that *Rpi-amr1* gene family diversity facilitates detection of diverse paralogs and alleles of the recognized effector, enabling broad-spectrum and durable resistance against *P. infestans*.

## Introduction

Potato is the fourth most important directly-consumed food crop world-wide^1^. *Phytophthora infestans*, an oomycete pathogen, causes late blight disease in potato, and can result in complete crop failure. Disease management is primarily based on repeated fungicide applications (10-25 times per season in Europe). However, fungicide-resistant races have emerged^2^.

To elevate late blight resistance, *Resistance* to *Phytophthora infestans* (*Rpi*) genes were identified in wild relatives of potato and used for resistance breeding^3^. More than 20 *Rpi* genes have been mapped and cloned from different *Solanum* species^4^. All encode coiled-coil (CC), nucleotide binding (NB), leucine-rich repeat (LRR) (NLR) proteins^5^ and some require helper NLR proteins of the NRC family^6^. However, most cloned *Rpi* genes have been broken by *P. infestans*^7^. Provision of durable late blight resistance for potato remains a major challenge.

NLR-mediated immunity upon effector recognition activates “effector-triggered immunity” (ETI)^8^. In oomycetes, all identified recognized effectors, or avirulence (*Avr*) genes, carry a signal peptide and an RxLR motif^9^. 563 RxLR effectors were predicted from the *P. infestans* genome, enabling identification of the recognized effectors^10,11^. Many *P. infestans* effectors show signatures of selection to evade recognition by corresponding NLR proteins^12^. NLR genes also show extensive allelic and presence/absence variation in wild plant populations^13,14^ and known *Resistance (R)* gene loci like *Mla, L, Pi9, RPP1* and *RPP13* from barley, flax, rice and Arabidopsis show substantial allelic polymorphism^15-18^. Remarkably, different *Mla* alleles can recognize sequence-unrelated effectors^19,20^.

Technical advances like RenSeq (Resistance gene enrichment and Sequencing) and PenSeq (Pathogen enrichment Sequencing) enable rapid definition of allelic variation and mapping of plant *NLRs*, or discovery of variation in pathogen effectors^21-23^. Combined with single-molecule real-time (SMRT) sequencing, SMRT RenSeq enabled cloning of *Rpi-amr3* from *Solanum americanum*^24^. Similarly, long read and cDNA PenSeq enabled us to identify *Avramr1* from *P. infestans*^25^.

In this study, we further explored the genetic diversity of *S. americanum*, and by applying sequence capture technologies, we fine-mapped and cloned *Rpi-amr1* from *S. americanum*, (usually) located on the short arm of chromosome 11. Multiple *Rpi-amr1* homologs were found in different *S. americanum* accessions and in relatives, including *Solanum nigrescens* and *Solanum nigrum*. Functional alleles show extensive allelic variation and confer strong, broad-spectrum resistance to all 19 tested diverse *P. infestans* isolates. Although differential recognition was found between different *Rpi-amr1* and *Avramr1* homologs, all *Rpi-amr1* alleles recognize the *Avramr1* homologs from *Phytophthora parasitica* and *Phytophthora cactorum*. Our study reveals unique properties of genetic variation of *R* genes from “non-host” species.

## Results

### *Rpi-amr1* maps to the short arm of chromosome 11

We previously investigated *S. americanum* and isolated *Rpi-amr3* from an accession 944750095 (SP1102)^24^. To discover new *Rpi-amr* genes, we characterized additional 14 lines of *P. infestans*-resistant *S. americanum* and close relatives *S. nigrescens* and *Solanum nodiflorum* by crossing them to a susceptible (S) *S. americanum* line 954750186 (hereafter SP2271) (Table 1). To avoid self-pollination, a resistant parent was always used as a pollen donor. All the corresponding F1 plants (6-10 per cross) were resistant in a detached leaf assay (DLA) (Table 1). Around 60-100 F2 progeny derived from each self-pollinated F1 plant were phenotyped by DLA using *P. infestans* isolate 88069^26^. The F2 progenies that derived from the resistant parents with working numbers SP1032, SP1034, SP1123, SP2272, SP2273, SP2360, SP3399, SP3400, SP3406, SP3408 and SP3409 segregated in a ratio suggesting the presence of a single (semi-) dominant resistance gene (fitting 3:1 or 2:1 [likely due to segregation distortion], R:S - resistant to susceptible - ratio). Two crosses showed a 15:1 segregation (resistant parent SP2300 and SP2307), suggesting the presence of two unlinked resistance genes, while SP1101 showed no susceptible plants in 100 individuals, suggesting the presence of three or more resistance genes.

**Table 1.**
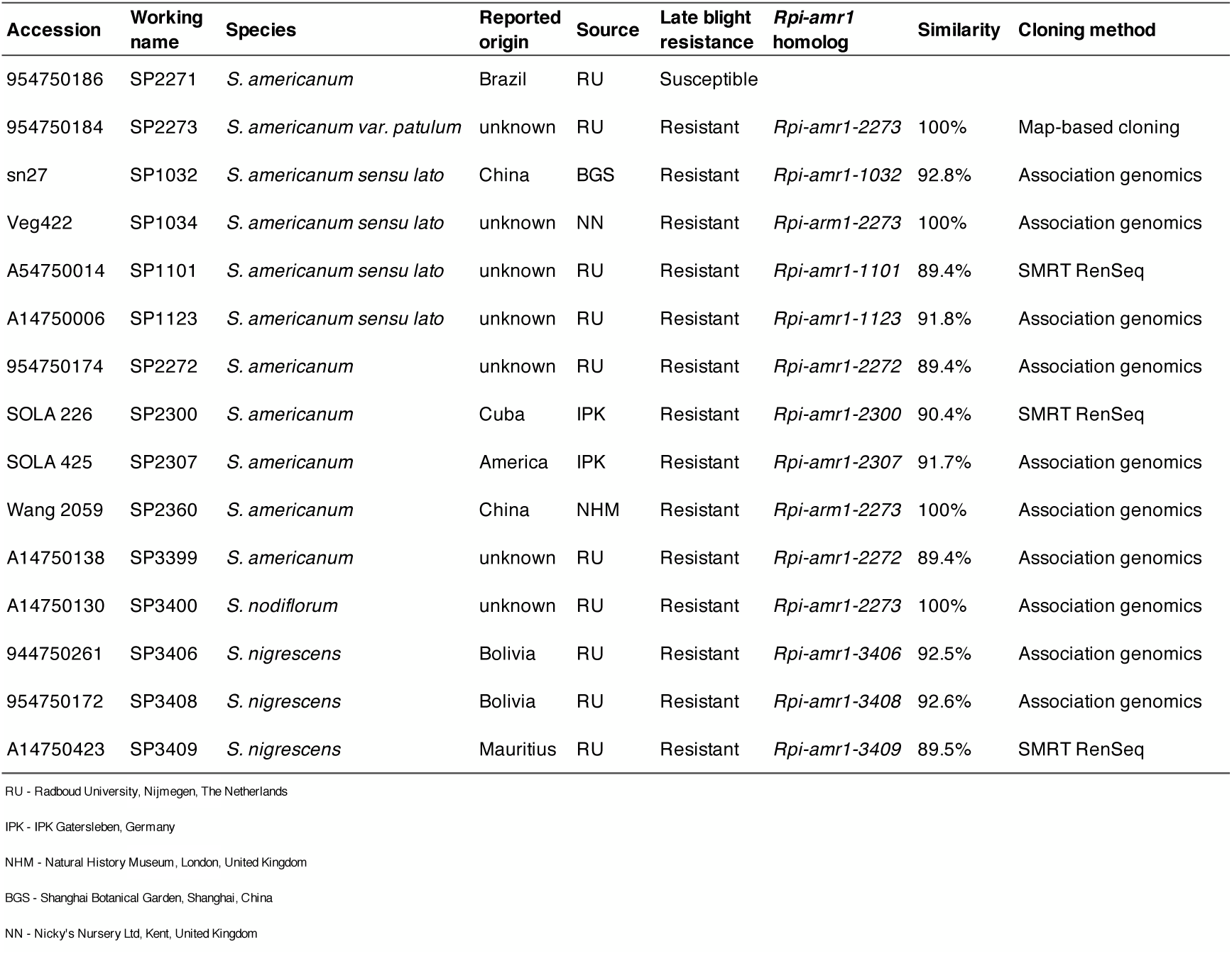
*S. americanum, S. nodiflorum* and *S. nigrescens* accessions used in this study and the corresponding *Rpi-amr1* homologs.

To identify *Rpi* genes from these resistant *S. americanum* accessions, we prioritized an F2 population derived from resistant parent SP2273 and named the corresponding gene *Rpi-amr1*. Using markers from RenSeq, genotyping by sequencing (RAD markers) and Whole Genome Shotgun sequencing (WGS), the *Rpi-amr1* gene was mapped in a small population (n=188 gametes) to the short arm of chromosome 11, between markers RAD_3 and WGS_1 (Fig. 1a, Table S1, S2). We expanded the mapping population and developed a PCR marker WGS_2 that co-segregated with resistance in 3,586 gametes (Fig. 1b, Table S2). To generate the physical map of the target interval from SP2273, a BAC library was generated. Two BAC clones (12H and 5G) covering the target interval were isolated and sequenced on the PacBio RSII platform, and assembled into a single contig of 212 kb (Fig. 1c). We predicted 11 potential coding sequences on the BAC_5G, nine of which encode *NLR* genes (Fig. 1c). These *NLR* genes belong to the CNL class and have 80-96% between-paralog identity.

To define which of these *NLR* genes are expressed, cDNA RenSeq data of the resistant parent SP2273 were generated and mapped to the BAC_5G sequence. Seven out of nine *NLR* genes were expressed. These genes - *Rpi-amr1a, b, c, d, e, g* and *h -* were tested as candidate genes for *Rpi-amr1* (Fig. 1c).

**Fig. 1.**
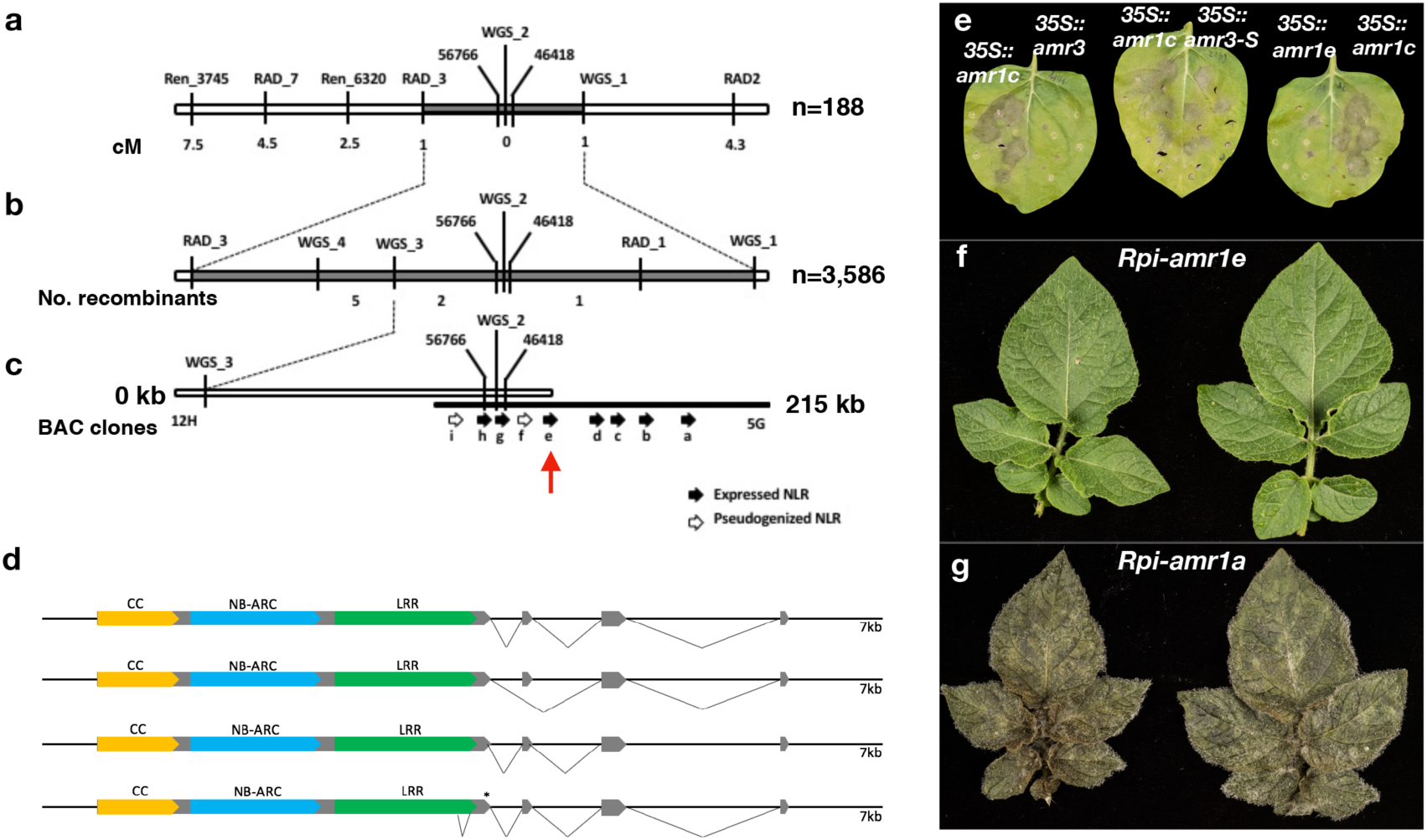
Map-based cloning of *Rpi-amr1* and its resistance to *P. infestans*. (a) Mapping of *Rpi-amr1* in a small F2 population (n=188 gametes); the names of the markers and genetic distances are shown above or below the bar. (b) Fine mapping of *Rpi-amr1* in the F2 population of 3,586 gametes. The names of the markers and the number of recombinants are shown above or below the bar. (c) Physical map of the target *Rpi-amr1* interval based on the assembled BAC contig. The markers present on the BAC are shown. The predicted NLR genes are depicted as black arrows (expressed NLRs) or empty arrows (pseudogenized NLRs). *Rpi-amr1* (formerly *Rpi-amr1e*) is indicated by a red arrow. (d) Four *Rpi-amr1* transcripts detected by 3’ RACE PCR. (e) Leaves of *N. benthamiana* plants were infiltrated with the binary vector pICSLUS0003∷*35S* overexpressing either the late blight resistance gene *Rpi-amr3* (positive control), one of seven *Rpi-amr1* candidates, or the non-functional *Rpi-amr3-S* (negative control). Leaves were inoculated with *P. infestans* strain 88069 24 h after infiltration. Only leaves infiltrated with *Rpi-amr3 and Rpi-amr1e* (pictured) showed reduced pathogen growth, whereas *P. infestans* grew well in the presence of the remaining *Rpi-amr1* candidates. Only *Rpi-amr1c* is shown as the phenotype of all other non-functional candidate genes was indistinguishable. Photographs were taken 9 dpi. (f) Transgenic potato cv. Maris Piper which expresses *Rpi-amr1* under the native regulatory elements is resistant to *P. infestans* isolate 88069 (top), displaying no symptoms at the spot of inoculation. Each leaflet was inoculated with a droplet containing approximately 1,000 zoospores; photographs were taken 9 dpi. (g) The control plants carrying the non-functional candidate *Rpi-amr1a* show large necrotic lesions and sporulation. Each leaflet was inoculated with a droplet containing approximately 1,000 zoospores; photographs were taken 9 dpi.

### *Rpi-amr1e* confers resistance in *Nicotiana benthamiana* and cultivated potato

To test the function of the seven candidate genes, we cloned their open reading frames into a binary expression vector under control of the 35S promoter. *Rpi-amr3* was used as a positive control and the non-functional *Rpi-amr3-S* was used as a negative control. The constructs carrying each of the seven candidate genes were transiently expressed after *Agrobacterium* infiltration into *N. benthamiana* leaves, which were subsequently inoculated with the *P. infestans* isolate 88069 as described^24^. *P. infestans* growth was observed six days post inoculation (dpi). Only *35S::Rpi-amr1e*-infiltrated leaves showed reduced pathogen growth at 6 dpi compared to other candidate genes like *Rpi-amr1c*, or negative control *Rpi-amr3-S*. (Fig. 1e). Hence, we conclude that *Rpi-amr1e* is the functional *Rpi-amr1* (hereafter) gene from *S. americanum* SP2273.

To test if *Rpi-amr1* confers late blight resistance in potato, we cloned it with its native promoter and terminator, and generated transgenic potato cultivar Maris Piper plants carrying *Rpi-amr1*. A non-functional paralog *Rpi-amr1a* was also transformed into Maris Piper as a negative control. As in the transient assay, stably transformed *Rpi-amr1* lines resisted *P. infestans* 88069 in potato (Fig. 1f), but *Rpi-amr1a*-transformed plants did not (Fig. 1g).

### *Rpi-amr1* is a four exon CC-NLR

To characterize the structure of *Rpi-amr1*, we mapped the cDNA RenSeq data to the full length *Rpi-amr1* gene, and found four alternatively spliced forms of *Rpi-amr1*. The most abundant form, supported by >80% of reads, comprises four exons encoding a protein of 1,013 amino acids. This was confirmed with 3’ RACE PCR (Fig. 1d). The Rpi-amr1 is a typical CC-NB-LRR resistance protein, with a coiled-coil domain (CC; amino acids 2-146), nucleotide binding domain (NB-ARC; amino acids 179-457) and leucine-rich repeats (LRR; located between amino acids 504-900) which are all positioned in the first exon (1-918 aa, Fig. 2a). The remaining three short exons (amino acids 919-943, 944-1002 and 1,003-1,013) lack homology to any known domains. No integrated domains^27^ were found in the Rpi-amr1 protein.

**Fig. 2.**
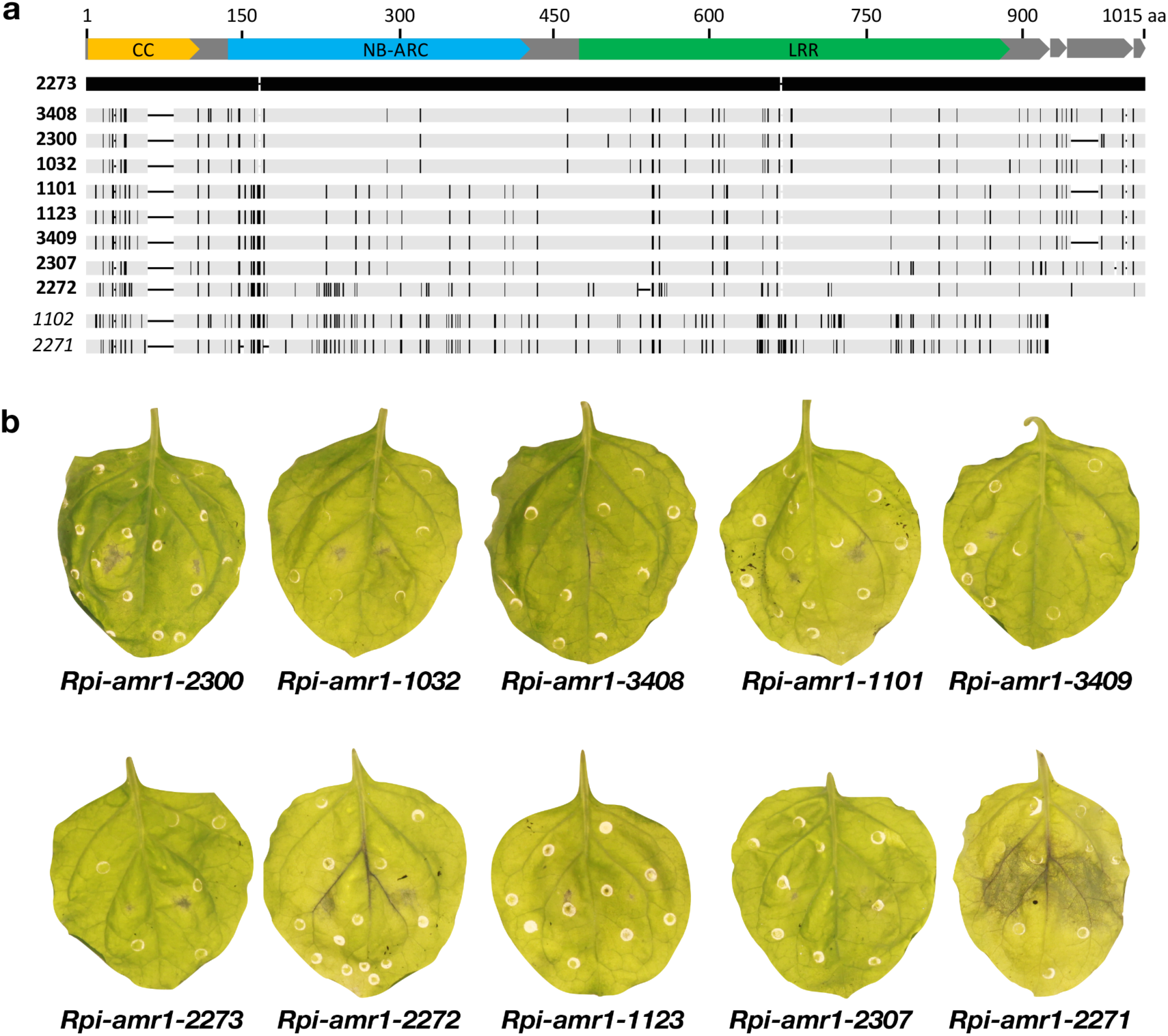
Schematic representation of amino acid sequence alignment of *Rpi-amr1* homologs (a) and *P. infestans* resistance in transient assay (b). (a) The exons and the conserved NLR domains are highlighted at the top of the alignment (exons, grey; CC, orange; NB-ARC, blue; LRR, green). Black bars in the alleles indicate the polymorphic nucleotides and indels as compared with *Rpi-amr1-2273*. The numbers next to the alleles refer to the accession working numbers (Table 1). Figure drawn to the scale. (b) Nine *Rpi-amr1* homologs provide resistance to *P. infestans* in transient complementation assay. *Rpi-amr1* genes with native regulatory elements were infiltrated into *N. benthamiana* leaves. At 1 dpi, leaves were cut off and drop inoculated with 10 µl of zoospore suspension (50,000/mL) from *P. infestans* isolate 88069. The non-functional *Rpi-amr1-2271* homolog from susceptible accession SP2271 was used as negative control. Photographs were taken 8 dpi.

### Functional *Rpi-amr1* homologs were identified from multiple resistant *S. americanum* and relatives

Previously, we found at least 14 *S. americanum* accessions and related species that resist late blight (Table 1). To test if *Rpi-amr1* contributes to late blight resistance in other resistant *S. americanum* accessions, we genotyped 10-50 susceptible F2 plants of the populations derived from resistant accessions, with *Rpi-amr1* linked markers (markers 3745 and 56766, Fig. 1 and Table S2). We found that in SP1032, SP1034, SP1123, SP2272, SP2307, SP2360, SP3399, SP3400, SP3406 and SP3408, resistance is linked to the *Rpi-amr1* locus. To test if in these accessions the resistance is conferred by functional *Rpi-amr1* homologs, we performed SMRT RenSeq-based *de novo* assembly of each resistant accession, and looked for homologs with the greatest identity to *Rpi-amr1*. For accessions SP2307, SP3399 and SP3406, we also used cDNA RenSeq to monitor their expression. We mapped *de novo* contigs to the coding sequence of *Rpi-amr1* allowing for 15% mismatches and gaps, and selected the closest homolog as a candidate *Rpi-amr1* ortholog (Table S3). In three resistant parents, namely SP1034, SP2360 and SP3400, the functional alleles showed 100% identity at the amino acid level to Rpi-amr1, while amino acid sequences from the remaining accessions had as little as 89% identity to the functional Rpi-amr1 (Table S3). As described previously, we transiently expressed the closest related candidate *Rpi-amr1* homologs in *N. benthamiana* leaves followed by DLA with *P. infestans* isolate 88069, and verified their functionality. The unique homologs of *Rpi-amr1-2273* were named as *Rpi-amr1-1032, Rpi-amr1-1123, Rpi-amr1-2272, Rpi-amr1-2307* and *Rpi-amr1-3408*.

For some accessions, like SP1101 and SP2300, the *Rpi-amr1*-linked markers gave ambiguous results, so we directly performed bulked segregant analysis (BSA) and RenSeq. Additional *Rpi-amr1* co-segregating paralogs, *Rpi-amr1-1101* and *Rpi-amr1-2300*, were identified and verified in transient assays as above (Fig. 2b).

Similarly, we inspected an F2 population derived from *S. nigrescens* accession SP3409 (Table 1). We applied BSA RenSeq and SMRT RenSeq to the resistant parents and F2 segregating population, and we found five candidate *NLRs* belonging to the same *Rpi-amr1* clade, all of which are expressed. The five candidates were cloned, and transient assays verified one of them as a functional *Rpi-amr1* homolog, *Rpi-amr1-3409*. However, *Rpi-amr1-3409* does not co-segregate with *Rpi-amr1-*linked markers. We used GenSeq sequence capture-based genotyping (Chen *et al*. 2018), and found that *Rpi-amr1-3409* locates on chromosome 1, based on the potato DM reference genome^28^. This result suggests that a fragment of DNA that locates on distal end of the short arm of chromosome 11 in other resistant accessions was translocated to the distal end of the long arm of chromosome 1 in SP3409.

When the full-length amino acid sequences of nine *Rpi-amr1* homologs were aligned, the polymorphisms between different functional alleles were found to be distributed through all domains including the LRR region (Fig. 2a and Fig. S1).

Taken together, by using BSA RenSeq, SMRT RenSeq, cDNA RenSeq, association genomics and GenSeq, we cloned eight additional functional *Rpi-amr1* homologs from different resistant accessions, of which all confer resistance to *P. infestans* 88069 in transient assays. The closest *Rpi-amr1* homolog from susceptible parent SP2271 does not confer resistance (Fig. 2b).

### *Rpi-amr1* is present in hexaploid *S. nigrum* accessions

Most *S. nigrum* accessions are highly resistant to *P. infestans* and *S. nigrum* has been reported to be a “non-host” to *P. infestans*^29^, even though rare accessions are susceptible^30^. *S. americanum* may be the diploid ancestor of hexaploid *S. nigrum*^31^. To test if *Rpi-amr1* also contributes to late blight resistance in *S. nigrum*, we amplified and sequenced the first exon of *Rpi-amr1* from four resistant and one reported susceptible *S. nigrum* accessions^30^. From three resistant accessions (SP1095, SP1088 and SP1097; Table S4), we amplified sequences with >99% nucleotide identity to *S. americanum Rpi-amr1-2273* (Fig. S2). *Rpi-amr1-1104* was more polymorphic, with 96.7% nucleotide identity to *Rpi-amr1-2273*, and primers used for allele mining did not amplify anything from the susceptible line SP999. These data suggest that *Rpi-amr1* homologs are present in some *S. nigrum* accessions and were most likely inherited from *S. americanum*.

### *Rpi-amr1* confers broad-spectrum late blight resistance in cultivated potato

To test the scope of late blight resistance conferred by *Rpi-amr1* and its homologs, we generated stably transformed transgenic potato cv Maris Piper plants carrying *Rpi-amr1-2272* and *Rpi-amr1-2273*, the most diverged of the homologs (Table S3), and inoculated them by DLA with 19 *P. infestans* isolates from UK, the Netherlands, Belgium, USA, Ecuador, Mexico and Korea (Table 2). Many of the tested *P. infestans* isolates can defeat multiple *Rpi* genes (Table 2). Our DLAs show that Maris Piper carrying *Rpi-amr1-2272* or *Rpi-amr1-2273* resist all 19 tested *P. infestans* isolates, while the wild-type Maris Piper control is susceptible to all of them. This indicates that *Rpi-amr1* confers broad-spectrum resistance against diverse *P. infestans* races.

**Table 2.** Phenotypes of potato plants stably transformed with *Rpi-amr1-2272* and *Rpi-amr1-2273* after inoculation with multiple isolates of *P. infestans*.

| Isolate | *Rpi-amr1-2272* | *Rpi-amr1-2273* | Maris Piper | Origin | Race<sup>e</sup> |
| --- | --- | --- | --- | --- | --- |
| NL00228 | R | R | S | The Netherlands | 1.2.4.7 |
| US23 | R | R | S | USA | n.a. |
| 3928A<sup>a</sup> | R | R | S | UK | 1.2.3.4.5.6.7.10.11<sup>f</sup> |
| EC3626<sup>b</sup> | R | R | S | Ecuador | n.a. |
| NL14538<sup>c</sup> | R | R | S | The Netherlands | n.a. |
| NR47UH<sup>d</sup> | R | R | S | UK | 1.3.4.7.10.11<sup>f</sup> |
| T30-4 | R | R | S | The Netherlands | n.a. |
| USA618 | R | R | S | USA | 1.2.3.6.7.10.11 |
| KPI15-10 | R | R | S | Korea | n.a. |
| IPO-C | R | R | S | Belgium | 1.2.3.4.5.6.7.10.11 |
| PIC99189 | R | R | S | Mexico | 1.2.5.7.10.11 |
| UK7824 | R | R | S | UK | n.a. |
| PIC99177 | R | R | S | Mexico | 1.2.3.4.7.9.11 |
| VK98014 | R | R | S | The Netherlands | 1.2.4.11 |
| NL08645 | R | R | S | The Netherlands | n.a. |
| PIC99183 | R | R | S | Mexico | 1.2.3.4.5.7.8.10.11 |
| NL11179 | R | R | S | The Netherlands | n.a. |
| EC1<sup>b</sup> | R | R | S | Ecuador | 1.3.4.7.10.11 |
| NL01096 | R | R | S | The Netherlands | 1.3.4.7.8.10.11 |

<sup>a</sup> Clonal lineage EU\_13\_A2, or “Blue13”
<sup>b</sup> Overcomes *Rpi-vnt1*
<sup>c</sup> Overcomes *Rpi-vnt1* and partially *Rpi-blb1*, *Rpi-blb2*
<sup>d</sup> Clonal lineage EU\_6\_A1, commonly known as “Pink6”
<sup>e</sup> Summarized in<sup>32</sup>
<sup>f</sup> See<sup>33</sup>

### Differential recognition by *Rpi-amr1* alleles of *Avramr1* homologs

*Avramr1* (*PITG_07569*) was identified in *P. infestans* race T30-4 by long-read and cDNA PenSeq, and multiple *Avramr1* homologs were identified in four *P. infestans* isolates and classified into four subclades^25^. To investigate if all nine cloned *Rpi-amr1* homologs could recognize diverse *Avramr1* homologs from different *P. infestans* isolates, in addition to *Avramr1* from race T30-4 that corresponds to clade A, we synthesized three *Avramr1* homologs *Avramr1-13B1, Avramr1-13C2* and *Avramr1-13D1* from isolate 3928A (EU_13_A2, commonly known as “Blue 13”), corresponding to clades B, C and D, respectively (Fig. 3). We also synthesized the *Avramr1* homologs from *P. parasitica* and *P. cactorum*^25^. These six *Avramr1* homologs were co-expressed in *N. benthamiana* by agro-infiltration in all possible combinations with nine functional *Rpi-amr1* homologs and the non-functional *Rpi-amr1-2271* as a negative control (Fig. 3).

**Fig. 3.**
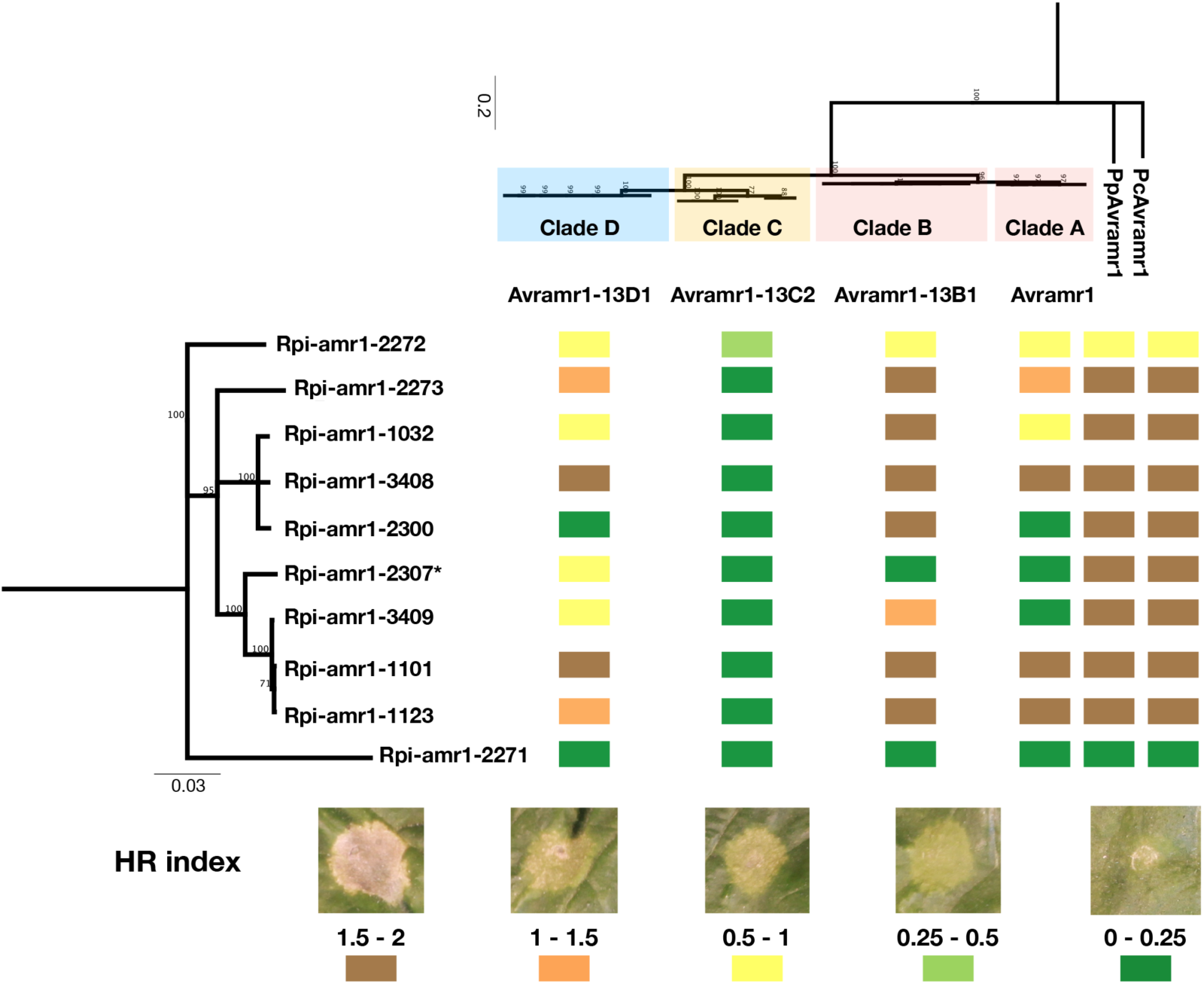
Differential recognition of *Rpi-amr1* and *Avramr1* homologs. Four *Avramr1* homologs representing clades A-D, and *P. parasitica* and *P. cactorum* homologs were co-infiltrated with ten Rpi-amr1 homologs, including a non-functional homolog *Rpi-amr1-2271*, into *N. benthamiana* leaves. Colours from green to brown represent the strength of HR scored from 0 to 2 (see bottom panel). N=3. Left: phylogenetic tree of nine functional *Rpi-amr1* homologs and non-functional homolog *Rpi-amr1-2271*. Top: phylogenetic tree of *Avramr1* homologs from four isolates of *P. infestans*. * Stable *Rpi-amr1-2307 N. benthamiana* transformants show HR upon transient expression of *Avramr1* and *Avramr1-13B*1.

We found that different combinations of *Rpi-amr1* alleles and *Avramr1* homologs led either to strong, weak or no HR phenotype in transient assay, but the non-functional *Rpi-amr1-2271* allele failed to recognize any *Avramr1* homologs (Fig. 3). *Rpi-amr1-2300* and *Rpi-amr1-2307* recognized one *Avramr1* homolog each, but others detected *Avramr1* homologs from more than one clade. Clade C, represented here by *Avramr1-13C2*, is usually not expressed^25^, and when expressed from 35S promoter, this effector was not recognized by most *Rpi-amr1* homologs, though a weak HR was observed upon co-expression with *Rpi-amr1-2272. Avramr1-13D1* belongs to Clade D, which is absent in T30-4 but present in four other sequenced isolates^25^, and was recognized by all but one (*Rpi-amr1-2300*) homologs in the transient assay. Surprisingly, two *Avramr1* homologs from *P. parasitica* and *P. cactorum* are strongly recognized by all functional *Rpi-amr1* homologs, apart from *Rpi-amr1-2272* which showed a weaker HR (Fig. 3).

Collectively, our data shows that *Rpi-amr1*/*Avramr1* homolog pairs provoke quantitatively and qualitatively different HRs, but all functional *Rpi-amr1* homologs detect at least one *Avramr1* homolog from *P. infestans* isolate 3928A.

### Both *Rpi-amr1*-mediated resistance and effector recognition are NRC2 or NRC3 dependent

We generated a phylogenetic tree for representative *Solanaceae* NLR proteins. Rpi-amr1 is grouped with clade CNL-3, from which no functional resistance genes were previously cloned (Fig. 4a). This phylogenetic affiliation suggested that Rpi-amr1 is likely to depend on the helper NRC clade because CNL-3 is among the large super-clade of NRC-dependent sensors (Fig. 4a)^6^.

**Fig. 4.**
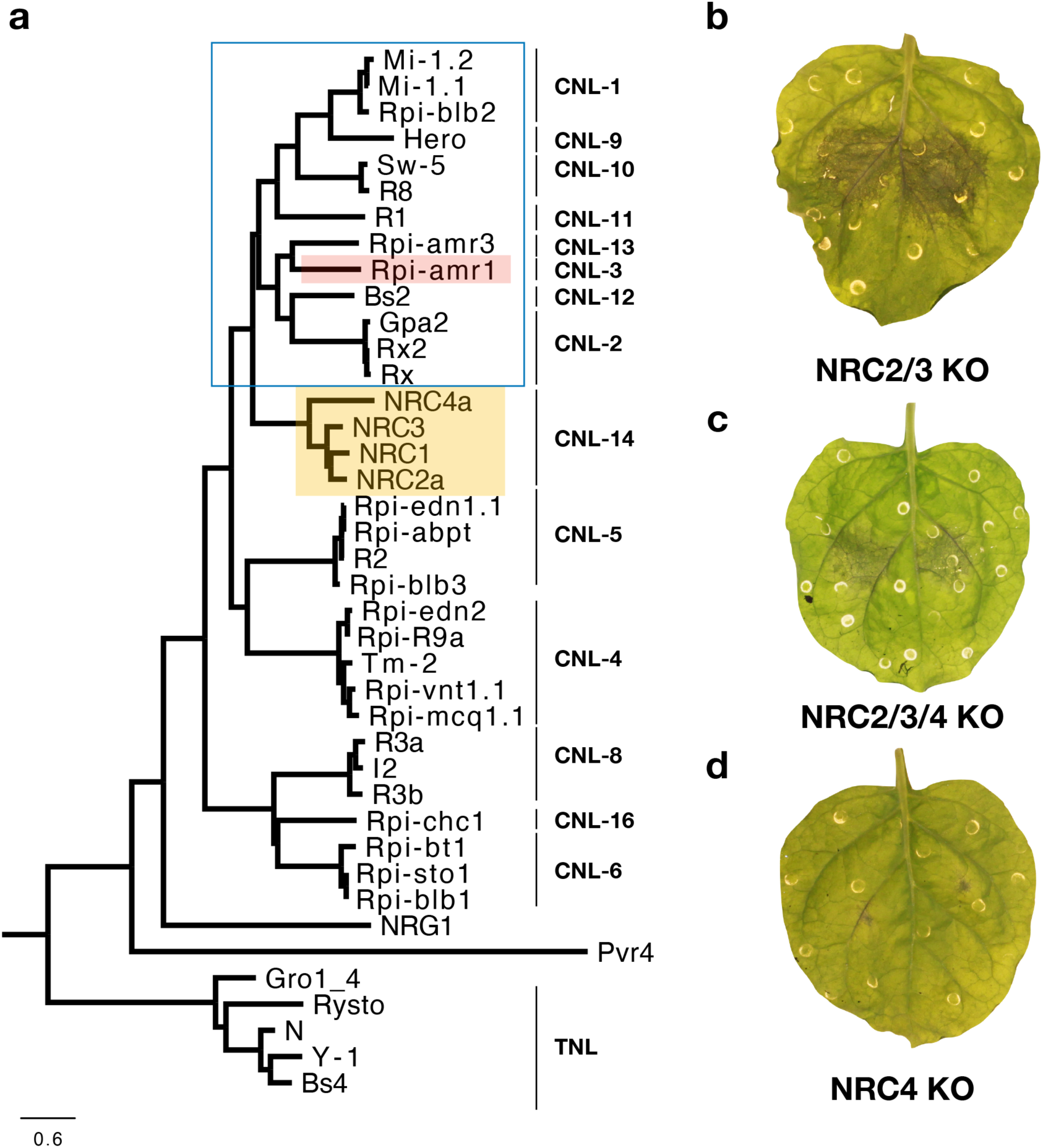
Rpi-amr1 is NRC2 or NRC3 dependent. (a) Phylogenetic analysis of Rpi-amr1 protein and other functional Solanaceae NLR proteins. The NLR clades shown here are as described previously^24^, the NRC-dependent sensor clades are marked by blue box. (b) Transient expression of *Rpi-amr1-2273* in *NRC2*/*NRC3* double knockout *N. benthamiana*, followed by zoospore inoculation of *P. infestans* isolate 88069, results in large necrotic lesions indicating lack of resistance. (c) Transient expression of *Rpi-amr1-2273* in *NRC2*/*NRC3*/*NRC4* triple knockout *N. benthamiana*, followed by zoospore inoculation of *P. infestans* isolate 88069, results in large necrotic lesions indicating lack of the resistance. (d) Transient expression of *Rpi-amr1-2273* in *NRC4* knockout *N. benthamiana*, followed by zoospore inoculation of *P. infestans* isolate 88069 results in small necrotic lesions indicating resistance.

To test this hypothesis, we transiently expressed *Rpi-amr1-2273* together with *PpAvramr1* in NRC4, NRC2/3 or NRC2/3/4 knock out *N. benthamiana* leaves^34,35^ (Fig. S3). The HR phenotype was abolished in NRC2/3 and NRC2/3/4 knockout plants (Fig. S4 c and b), but not in NRC4 knock-out or wild-type plants (Fig. S4 d and a). The HR was recovered when NRC2 or NRC3 was co-expressed in the NRC2/3/4 or NRC2/3 knock out plants, but co-expression of NRC4 did not complement the loss of HR phenotype in NRC2/3/4 knockout plants. (Fig. S4 b and c). We further showed that also *Rpi-amr1* mediated resistance is dependent on NRC2 or NRC3 but not NRC4, as transient expression of *Rpi-amr1-2273* followed by *P. infestans* infection restricted pathogen growth only in NRC4 knockout *N. benthamiana* plants (Fig. 4b). These data indicate that both the effector recognition and resistance conferred by *Rpi-amr1* is NRC2 or NRC3 dependent.

### High allelic diversity at *Rpi-amr1* was generated through inter-paralog and ortholog sequence exchange

*Rpi-amr1* alleles show relatively high nucleotide diversity (π=0.04), which could be an indication of balancing or diversifying selection (Table S5). In addition, *Rpi-amr1* alleles differ in their recognition of the *Avramr1* homologs (Fig. 3) which is also consistent with selection in a host-parasite co-evolutionary arms race. To test the hypothesis that allelic polymorphism at *Rpi-amr1* results from diversifying selection, we calculated diversity statistics and performed a McDonald-Kreitman test on both *Rpi-amr1* alleles and *Avramr1* homologs. As expected, *Avramr1* homologs show a signature consistent with balancing selection (Tajima’s D = 2.27) (Table S5). Remarkably, despite the high nucleotide diversity, no clear signals of balancing or diversifying selection were detected for *Rpi-amr1* (Tajima’s D = 0.09083) (Table S5). Aligning the *Rpi-amr1* alleles against the reference and scrutinizing the sequences in more detail provided further insights. The nucleotide similarity of alleles varies markedly across the *Rpi-amr1* homologs (Fig. 2a and Table S3); this pattern is consistent with occasional recombination between highly diverged alleles or paralogs.

To test whether recombination could explain the observed polymorphisms in *Rpi-amr1* alleles, we predicted the possible recombination events using 3SEQ. Several recombination events were detected between *Rpi-amr1* orthologs from different *S. americanum* accessions, and *Rpi-amr1* paralogs from SP2273 (Table S6). Some sequence exchanges were visualized using HybridCheck (Fig. S5)^36^, and these data suggest that sequence exchange occurred between functional *Rpi-amr1* alleles and paralogs. To confirm these findings, we mapped all cloned *Rpi-amr1* CDS back to the BAC_5G sequence from accession SP2273 (Fig. S6). As expected, some *Rpi-amr1* homologs (e.g. SP2300 and SP2272) show a perfect match with the fourth NLR, and show a distribution of high identity that reflects the intron-exon structure. For some homologs (e.g. 2271), 5’ end sequences match different NLR sequences on the BAC_5G and for others (e.g. 2275) part of the sequence is highly diverged from BAC_5G. Taken together, our results indicate that the polymorphism of *Rpi-amr1* alleles appears to have arisen partly due to sequence exchange between highly diverged alleles and paralogs, and not just through mutation accumulation.

## Discussion

Achieving durable resistance is the ultimate goal of resistance breeding. Here, we report significant progress towards durable resistance against potato late blight. Most cloned late blight resistance genes derive from wild potatoes, and many have been overcome by one or more *P. infestans* strains^37^. Conceivably, resistance to *P. infestans* in nearly all *S. americanum* and *S. nigrum* accessions is due to multiple *NLR* genes, as zoospores from *P. infestans* can germinate on *S. nigrum* leaves but penetration is stopped by strong HR^29,38^. *Rpi* genes from plant species that only rarely support pathogen growth have likely not participated, or are no longer participating, in an evolutionary arms race with *P. infestans*, and hence, the pathogen’s effectors have not (yet) evolved to evade detection by these *Rpi* genes. Under this scenario, a pre-existing standing variation in the pathogen for overcoming such *Rpi* genes is either absent or extremely rare. This makes such genes promising candidates for provision of broad-spectrum and durable late blight resistance, provided they are not deployed alone which facilitates one-step genetic changes in the pathogen to evade them, but rather in combination with other genes, as in the source plant^39^.

We report here a novel, broad-spectrum *S. americanum* resistance gene, *Rpi-amr1*. We also identified eight additional *Rpi-amr1* alleles from different *S. americanum* accessions and relatives, including one *Rpi-amr1* allele that translocated to the long arm of chromosome 1. Allele mining also suggested the presence of *Rpi-amr1* homologs in *S. nigrum*. All nine cloned *Rpi-amr1* alleles confer late blight resistance in transient assays in *N. benthamiana*, and both *Rpi-amr1-*2272 and *Rpi-amr1-2273* in potato cv Maris Piper background confer resistance to all 19 tested *P. infestans* isolates from different countries, many of which overcome other *Rpi* genes. Thus, *Rpi-amr1* is widely distributed in germplasm of *S. americanum*, its relatives and *S. nigrum*, and may contribute to the resistance of nearly all accessions to *P. infestans*.

Many plant *R* genes and their corresponding *Avr* genes evolved differential recognition specificities with extensive allelic series for both *R* gene and *Avr* genes. Examples include *ATR1* and *RPP1* or *ATR13* and *RPP13* from *Hyaloperonospora arabidopsidis* and *Arabidopsis*^9^, *Avr567* and *L* genes from the rust *Melampsora lini* and flax^40^, and multiple and diverse recognized effectors from barley powdery mildew and *Mla* from barley. In the same manner, *Avramr1* and its homologs from several *P. infestans* races^25^ were found to be differentially recognized by high allelic variation at the *Rpi-amr1* gene. Remarkably though, the nucleotide diversity of the *R* gene did not show any of the hallmarks of diversifying or balancing selection.

Rather than through mutation accumulation, the high allelic variation observed at *Rpi-amr1* appears to have been generated partly by recombination between significantly diverged alleles and paralogs. The recombination events are likely to be rare relative to the mutation rate, given that the alleles carry many polymorphisms. This evolutionary scenario can explain the observed mosaic-like structure of high and low sequence similarities when the *Rpi-amr1* alleles were mapped against the contig based on two overlapping BAC clones. The deep coalescence of alleles that is implicit in this scenario can be generated by balancing selection, but we did not find evidence of such selection when analysing the nucleotide substitution patterns. Recombination between *Rpi-amr1* alleles could have eroded this signature of selection, as has been observed also in *Rp1* resistance genes in grasses^41^ and in the vertebrate immune genes of the major histocompatibility complex (MHC)^42,43^. Nucleotide sequence diversity across the *Rpi-amr1* alleles is correlated with only slight differences in *Avramr1* recognition specificity. *Rpi-amr1* alleles can even recognize multiple *Avramr1* paralogs from a single *P. infestans* strain, a scenario that might elevate durability of resistance. Since the *S. americanum* population recognizes multiple *Avramr1* alleles and paralogs, small mutational changes in *Avramr1* gene are unlikely to suffice to escape detection, which makes resistance-breaking less likely, thus promoting evolutionary durability of *Rpi-amr1*. We hypothesise that this enhanced recognition capacity could be key to the evolution of “non-host” resistance, offering an escape from the coevolutionary arms race. Conceivably, stacking *Rpi-amr1* alleles *in cis* could extend the recognition specificities, which could potentially lead to even more durable late blight resistance.

Intriguingly, two *Avramr1* homologs from *P. parasitica* and *P. cactorum* are recognized by all *Rpi-amr1* homologs. Presumably, these genes have been under even less selection pressure to evade *Rpi-amr1* recognition. This result indicates that *Rpi-amr1* has the potential to provide “non-host” type resistance in *S. americanum* against multiple oomycete pathogens like *P. parasitica* and *P. cactorum*, which can infect a wide range of hosts. As both the resistance and effector recognition of *Rpi-amr1* are *NRC2* or *NRC3* dependent, co-expression of *NRC2* or *NRC3* with *Rpi-amr1* might enable it to confer resistance to other *Phytophthora* species outside the *Solanaceae*.

In summary, we cloned *Rpi-amr1*, a broad-spectrum *Rpi* gene that contributes to the strong resistance of nearly all *S. americanum* accessions to late blight. The apparent redundancy across the *Rpi-amr1* gene family may serve an evolutionary function by broadening the scope for recognizing multiple *Avramr1* alleles and paralogs, and potentially reducing the probability of evolution of resistance-breaking strains. Stacking this type of *Rpi* gene with additional *Rpi* genes might help to turn host plants such as potato into non-hosts for late blight, enabling broad-spectrum and durable resistance.

## Methods

Methods and associated references are in supplementary information.

## Accession codes

Supporting raw reads and annotated BAC sequences were deposited in European Nucleotide Archive (ENA) under project number PRJEB38240.

## Acknowledgements

This research was financed from BBSRC grant BB/P021646/1 and the Gatsby Charitable Foundation. This research was supported in part by the NBI Computing infrastructure for Science (CiS) group through the provision of a High-Performance Computing Cluster. We would like to thank TSL bioinformatics team, transformation team and horticultural team for their support. We thank Experimental Garden and Genebank of Radboud University, Nijmegen, he Netherlands, IPK Gatersleben, Germany and Sandra Knapp (Natural History Museum, ondon, UK) for access to *S. americanum, S. nigrescens* and *S. nigrum* genetic diversity, and eert Kessel, Francine Govers and Paul Birch for providing *P. infestans* isolates.

## Author contributions

K.W., X.L., F.J., R.S., C.O. and J.D.G.J. designed the study. K.W., X.L., H.S.K., F.J., A.I.W., S.B., W.B., L.T. and T.S., performed the experiments. K.W., X.L., H.S.K., F.J., A.I.W., B.S., R.S., C.O., S.F., and J.M.C. analysed the data. K.W., X.L., H.S.K., F.J. and J.D.G.J. wrote the anuscript with input from all authors. V.G.A.A.V., B.B.H.W, C.-H.W., H.A. and S.K. contributed resources. K.W., X.L and H.S.K. made equivalent contributions and should be considered joint first authors. All authors approved the manuscript.

## Supplementary files

**Fig. S1:** Alignment of Rpi-amr1 proteins, including non-functional homolog from SP2271.

**Fig. S2:** Alignment of *Rpi-amr1-2273* and *Rpi-amr1* DNA sequences from *S. nigrum*.

**Fig. S3:** Genotypes and phenotypes of *NRC2/3* knockout *N. benthamiana*. (a). Amplicon sequencing results of the *NRC* loci of the *NRC2/3* knockout *N. benthamiana* line *nrc23*_1.3.1. sequences of the sgRNAs are marked in red; (b). The *NRC2/3* knockout line (*nrc23*_1.3.1) did not exhibit any growth defects when compared to the wild type plants. Six-week-old wild type and *NRC2/3* knockout *N. benthamiana* lines were used in the photograph.

**Fig. S4:** The effector recognition of *Rpi-amr1* is NRC2 or NRC3 dependent. The *Rpi-amr1 a*nd *Pp-Avramr1* were co-expressed by agro-infiltration on (a) wild type *N. benthamiana*; (b) RC2/3/4 knockout line; (c) NRC2/3 knockout line and (d) NRC4 knockout line. *NRC2, NRC3 o*r *NRC4* were co-expressed with *Rpi-amr1* or *Pp-Avramr1* on different knockout lines. *Rpi-amr1-2273* or *Avramr1* alone were used as negative controls.

**Fig. S5:** Sequence exchange between *Rpi-amr1* homologs. Sequence exchange events were visually checked and highlighted (b and d) or identified by HybridCheck (a and c). For HybridCheck, sequence similarity was visualised using the colours of an RGB colour triangle (top); deviation from the default red, green and blue at positions with the same colour indicates regions where two sequences share the same polymorphisms, which is indicative of intra- or inter-locus recombination. Line plot shows the percentage of SNPs shared at informative sites between sequences in each of the three pairwise combinations for the triplet.

**Fig. S6:** Cloned *Rpi-amr1* CDS were mapped back to BAC_5G using BLAT and visualized on the BAC sequence using the Sushi package.

**Table S1:** Linked RAD markers identified based on tomato reference genome.

**Table S2:** Molecular markers used in this study.

**Table S3:** Amino acid sequence similarity between Rpi-amr1 homologs.

**Table S4:** *S. nigrum* accessions used in this study.

**Table S5**: Tajima’s D analysis of *Rpi-amr1* and *Avramr1* homologs.

**Table S6**: Evidence of sequence exchange between *Rpi-amr1* orthologs and paralogs from P2273 using 3SEQ.

## Supplementary Materials and Methods

### Development of mapping populations

14 *P. infestans* resistant diploid *Solanum americanum* and relatives were used in this study Table 1). The F1 populations were generated by crossing with a susceptible *Solanum americanum* accession 954750186 (working name SP2271) as a female parent. Heterozygous F1 progeny was allowed to self-pollinate to generate F2 segregating populations, or further back-crossed to the susceptible parent and allowed to self-pollinate until resistance to *P. infestans* co-segregated as a monogenic trait.

### *P. infestans* infection assay

*P. infestans* isolates were cultured on rye and sucrose agar (RSA) medium at 18 °C for 10 days. Sporangia were washed off with cold water and incubated at 4°C for 1-2 h to induce zoospore release. Detached leaves were inoculated on the abaxial side with 10 µl droplets of zoospore suspension (50-100,000 per ml). The inoculated leaves were incubated at 18°C in high humidity under 16 h day/8 h night photoperiod conditions. Disease scoring was done at 5-7 ays after infection.

### DNA and RNA extraction

RenSeq experiments (both short- and long-reads protocols) were conducted on gDNA freshly extracted from young leaves using the DNeasy Plant Mini Kit (Qiagen) according to the manufacturer’s protocol. For the cDNA RenSeq experiment, RNA was extracted using TRI-Reagent (Sigma-Aldrich, MO, USA) and Direct-zol RNA MiniPrep Kit (Zymo Research, CA, USA), following manufacturer’s recommendations.

### Mapping of *Rpi-amr1*

To map the underlying resistance gene from the resistant parent 954750184 (working name SP2273), we generated an F2 segregating population which was phenotyped with *P. infestans i*solates EC1_3626 and 06_3928A. Selected resistant plants were self-pollinated and up to 100 plants from F3 populations were screened for resistance and susceptibility with *P. infestans i*solates EC1_3626 and 06_3928A. gDNA from susceptible F2 and F3 plants (BS pool), as well as gDNA from the resistant (R) and susceptible parent (S) were subjected to RenSeq using Solanaceae bait library^1^ and sequenced with Illumina GAII 76 bp paired-end reads. Pre-processing, assembly, mapping and SNP calling was performed as described earlier^1,2^.

The same gDNA samples were used in a RAD-seq experiment using PstI digestion and Illumina HiSeq sequencing, which was outsourced to Floragenex Inc. (OR, USA). Bioinformatic analysis was also performed by Floragenex using *Solanum lycopersicum* genome as a reference^3^. SNP calling resulted in sixteen polymorphic sites with eleven of them locating at the top of chromosome 11 (Supplementary table 1). The remaining ones were randomly distributed on chromosomes 4 and 1.

We additionally outsourced Whole Genome Shotgun sequencing (WGS) of R and S samples to BGI (BGI, Shenzhen, China) for ∼30 deep Illumina HiSeq sequencing with 100PE. Reads from the resistant parent were assembled as described in^2^ and we used our previously published i*n silico* trait mapping pipelines to perform SNP calling and detection of polymorphisms linked to disease resistance^1,2^. Contigs polymorphic between R and S parents were further aligned to the DM reference genome to identify their position.

Screening a set of markers derived from these three approaches on gDNA of 94 susceptible F2 and F3 plants identified 12 markers linked to resistance response that flank the *R* locus between 7.5 cM to one side and 4.3 cM to the other side (WGS, Table S1). Four of these markers were found to co-segregate with the resistance, and two others located around 1 cM on either side, CAPS marker RAD_3 to the distal side and the PCR marker WGS_1 to the proximal side (Figure 1). Both 1 cM markers were subsequently used to genotype 1,793 F2 plants, and we identified 228 recombinants (118 homozygous susceptible to one side and heterozygous to the other, 110 homozygous resistant to one side and heterozygous to the other).

The 118 informative recombinants (homozygous susceptible / heterozygous) were further genotyped using eight linked markers (Figure 1b), and tested in detached leaf assays for their response to *P. infestans* isolates EC1_3626 and 06_3928A. This revealed that markers CLC_3 (WGS_3) and RAD1 are flanking with a single recombination event for each marker, and CLC_2 (WGS_2), 56766 and 46418 are co-segregating with the resistance locus (Figure 1b). comparison of the linkage map (Figure 1) with the potato reference genome^4^ identified the homogeneous CNL-3 NLR gene sub-family to be within the co-segregating locus. This cluster comprises 18 members on potato reference chromosome 11.

### BAC clones identification and analysis

Construction and screening of 5x BAC library from resistant parent SP2273 was outsourced to BioS&T company (Quebec, Canada). Two candidate BAC clones (5G and 12H) were identified in PCR screen with WGS_2 marker-specific primers. BAC sequencing with RSII PacBio platform and bioinformatic analysis was outsourced to Earlham Institute (Norwich, UK); both BACs were assembled into single contigs with length of 125,327 bp (5G) and 144,006 bp (12H). While the co-segregating marker WGS_2 was present on both BAC clones, a further co-segregating marker WGS_3 was only present on 12H. The BACs were further assembled into one 212,773 contig (available in ENA under study number PRJEB38240). NLRs on the contig sequence were annotated using NLR-annotator^5^ and Geneious 8.1.2 build-in ORF prediction tool. Gene models were annotated manually using cDNA RenSeq data generated from *S. americanum* accession SP2273 as described below.

### 3’ RACE

Total RNA was extracted using RNeasy Plant Mini Kit (Qiagen) and treated with RNase-Free DNase (Qiagen) following manufacturer’s instructions. First strand cDNA was synthesized from total RNA using SuperScript™ First-Strand Synthesis System for RT-PCR (Invitrogen, CA, USA) with P7-oligoDT primer. The resulting product was amplified with P7- and gene specific primers by using KAPA HiFi HotStart ReadyMix PCR Kit (Kapa Biosystems, Cape Town, SA) and cloned into pCR™-Blunt II-TOPO vector by using Zero Blunt^®^ TOPO^®^ PCR Cloning Kit (Invitrogen) and transformation was performed using One Shot™ TOP10 Chemically Competent *E. coli* (Invitrogen). Isolation of plasmid DNA was performed with NucleoSpin^®^ Plasmid kit (MACHEREY-NAGEL, Duren, Germany).

### Resistance gene enrichment sequencing (RenSeq) and Gene enrichment sequencing GenSeq)

SMRT RenSeq, short-read RenSeq and cDNA RenSeq were performed as described previously^2^ and enriched libraries were sequenced at Earlham Institute, Norwich, UK (PacBio RSII, Illumina MiSeq) and Novogene, Hong Kong (Illumina HiSeq).

Illumina GenSeq was performed as described above (Illumina RenSeq), except GenSeq baits^6^ were used instead of RenSeq baits.

PacBio reads were processed and assembled using Geneious R8.1.8^7^ as described^2^. NLR coding sequences were predicted with Geneious and AUGUSTUS^8^ and annotated with NLR-parser^5^.

To infer linked polymorphisms, the quality control for Illumina paired-end reads was performed using Trimmomatic^9^ with standard settings. For the RenSeq, the paired reads were mapped to PacBio-assembled contigs from the resistant parent, while GenSeq reads were mapped to the reference DM genome (PGSC_DM_v4.03_pseudomolecules.fasta), using BWA mapper^10^ with default settings. PCR duplicates and unmapped reads were removed and Mpileup files to find out potential linked SNPs were created using SAMtools^11^. Mpileup files were processed with VarScan^12^ set to minimum read depth 20, minimum variant allele frequency threshold 0.1, and minimum frequency to call homozygote 0.98. The candidate SNPs were manually inspected using Savant genome browser^13^. TopHat^14^ with default settings was used to map cDNA Illumina reads to assembled PacBio data. All the tools used in this study were embedded in The Sainsbury Laboratory (TSL) customized Galaxy instance, if not stated otherwise.

### Transient complementation of a candidate genes in *N. benthamiana*

The candidate genes were PCR amplified from gDNA with their own promoters (1-2 kb upstream of start codon) and up to 1 kb terminator elements, and cloned into USER vector as described^2^. Transient complementation assays followed by *P. infestans* inoculation were performed as described^2^.

### Stable transformation of susceptible potato cultivar Maris Piper

Stable transgenic plants with constructs carrying *Rpi-amr1-2272, Rpi-amr1-2273* or *Rpi-amr1a* under the control of their native regulatory elements were created in the background of potato cultivar Maris Piper as described previously^15^. At least 10 independent transgenic lines were generated for each construct and tested for the presence of the transgene using gene specific primers. All positive *Rpi-amr1-2272* and *Rpi-amr1-2273* lines showed resistance in DLA with *P. infestans* isolate 88069, while *Rpi-amr1a* transgenic plants were fully susceptible. Selected lines of *Rpi-amr1-2272* and *Rpi-amr1-2273* were tested in DLA with 19 additional *P. infestans* isolates (Table 2). WT Maris Piper plants were used as a negative control.

### Generation of *NRC2/3* knockout *N. benthamiana*

*NRC4* and *NRC2*/*NRC3*/*NRC4* knockout *N. benthamiana* lines were described previously^16,17^. Knocking out of *NRC2*/*NRC3* in *N. benthamiana* were performed according to the methods described previously^16^. Forward primers CHW_sgNbNRCs and reverse primer JC_sgrna_R^16^ were used to clone sgRNA2.1-4, sgRNA3.1-4 into Golden Gate level 1 vectors for different positions. Constructs of sgRNAs targeting *N. benthamiana NRC2* and *NRC3* were assembled into level 2 vector pICSL4723 together with pICSL11017 (pICH47732::NOSp::BAR, Addgene no. 51145) and pICH47742::35S::Cas9^18^. Leaf discs of *N. benthamiana* were transformed with the binary vector pICSL4723 containing the BAR selection marker gene, Cas9 expression cassette, and sgRNAs targeting *NRC2* and *NRC3*. T0 transgenic plants were selected in the medium with phosphinothricin (2 mg/L) and then transferred into the soil. The progeny of the transformants were genotyped using amplicon sequencing as described previously^16^ (Fig. S6a). T3 populations from the selected T2 plants were used for further experiments. *NRC2/3* knockout line (*nrc23*_1.3.1) did not exhibit any growth defects when compared to the wild type plants (Fig. S6b).

### Phylogenetic tree construction

Phylogenetic tree was generated from protein sequences of the cloned *Solanaceae R* genes obtained from NCBI. Full-length sequences were aligned using ClustalW 1.74^19^ and the alignments were imported to the MEGA7^20^ to build a maximum-likelihood phylogenetic tree with Jones-Taylor-Thornton (JTT) substitution model and 100 bootstraps.

### Evolutionary analyses of *Rpi-amr1* and *Avramr1* homologs

CDS were aligned using MUSCLE^21^ as implemented in seaview^22^ with and without outgroup (the closest homologs from *S. lycopersicum* and *P. capsici* for *Rpi-amr1* and *Avramr1*, respectively). Calculations of diversity statistics and the MacDonald-Kreitmann Test were executed through DNAsp5.0^23^; DAMBE^24^ was used to rule out saturation. For *Rpi-amr1* homologs, the calculations were preformed separately on annotated full-length sequences as well as the individual domains.

We used 3SEQ^25^ to identify break points in the aligned CDS. To confirm gene conversion events in *Rpi-amr1*, we mapped the CDS back to the BAC_5G sequence using BLAT (minScore 1500, minMatch 93)^26^. The resulting .psl files were converted into .bed files using a custom R script, prior to visualization using the R package Sushi^27^.

### HybridCheck

For each accession, FASTA files of all *Rpi-amr1e* orthologs or *Rpi-amr1* paralogs in combinations of three (triplets) were generated and aligned using MUSCLE v3.8.31^21^. The sequence triplets were analysed using HybridCheck^28^ to detect and date recombination blocks between *Rpi-amr1* orthologs (sliding windows = 200bp) or paralogs (sliding windows = 100 bp); non-informative sites were removed from the sequence triplets. Figures showing sequence similarity were plotted (MosaicScale = 50) with HybridCheck and formatted using R v3.2.0 (https://www.r-project.org). The colour of each sequence window was calculated based on the proportion of SNPs shared between pairwise sequences at informative sites.

