## Supplementary figures and images for "A complex resistance locus in *Solanum americanum* recognizes a conserved *Phytophthora* effector"

### Fig S2

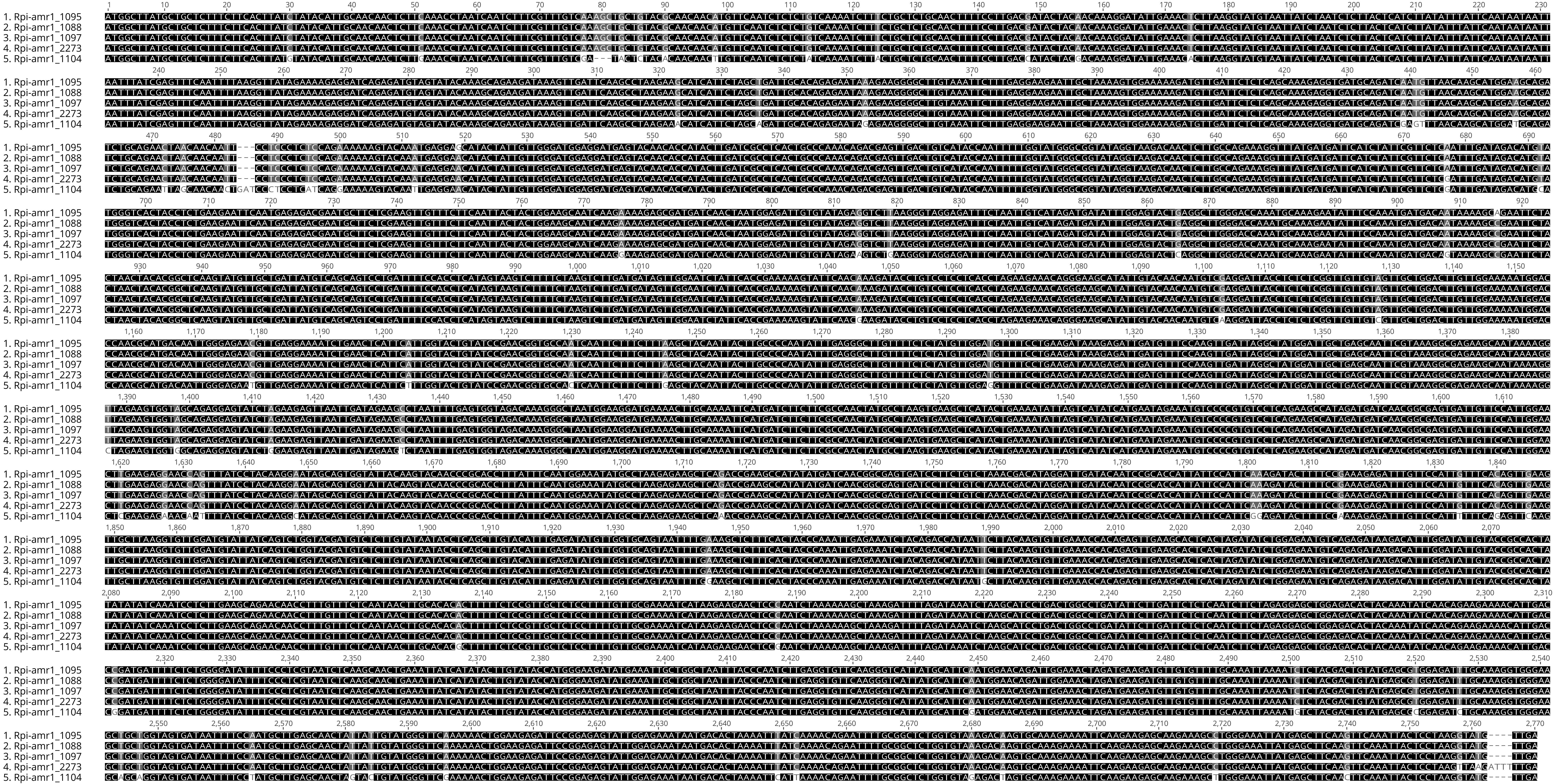

### Fig. S1

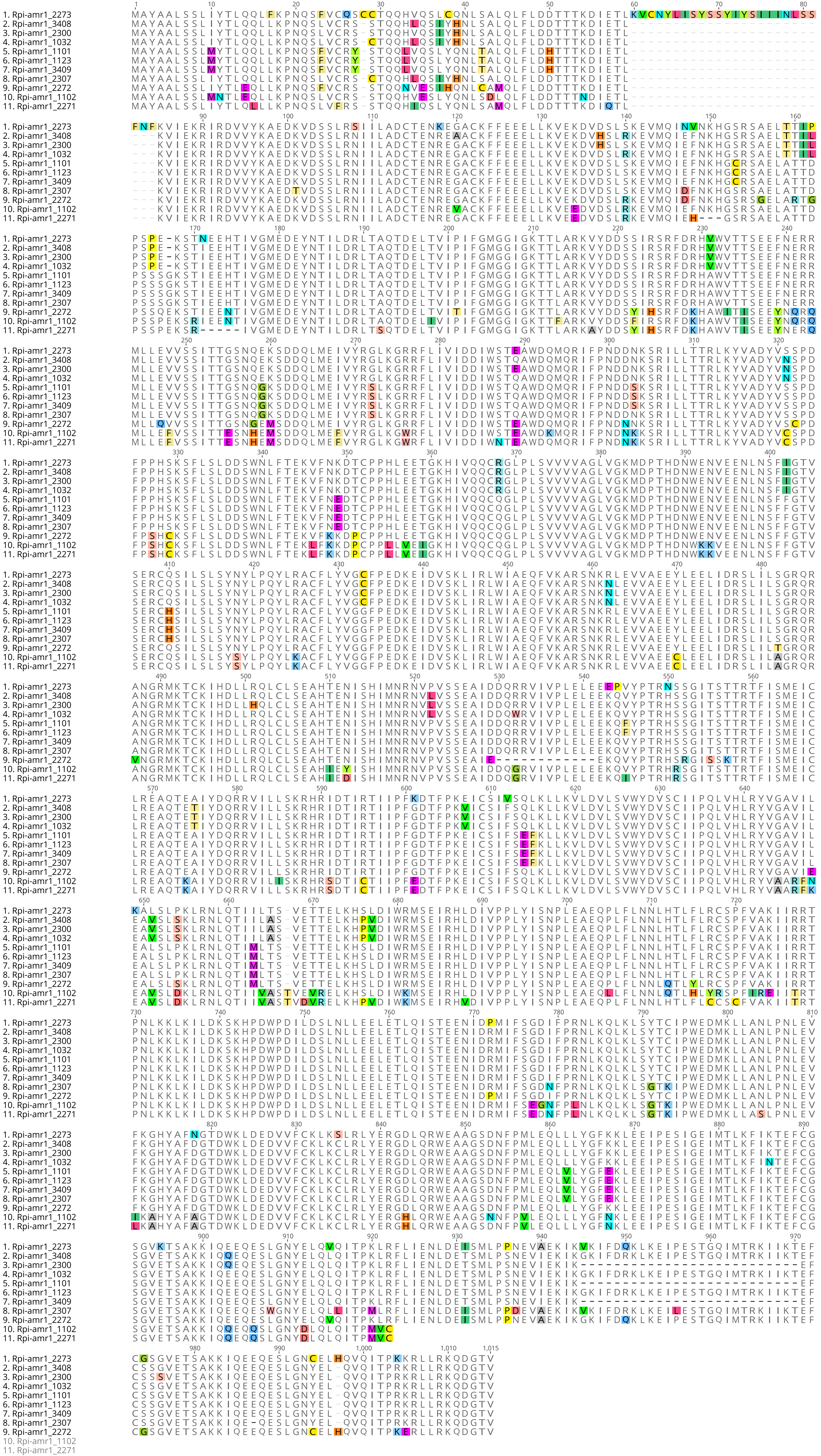

### Fig. S3

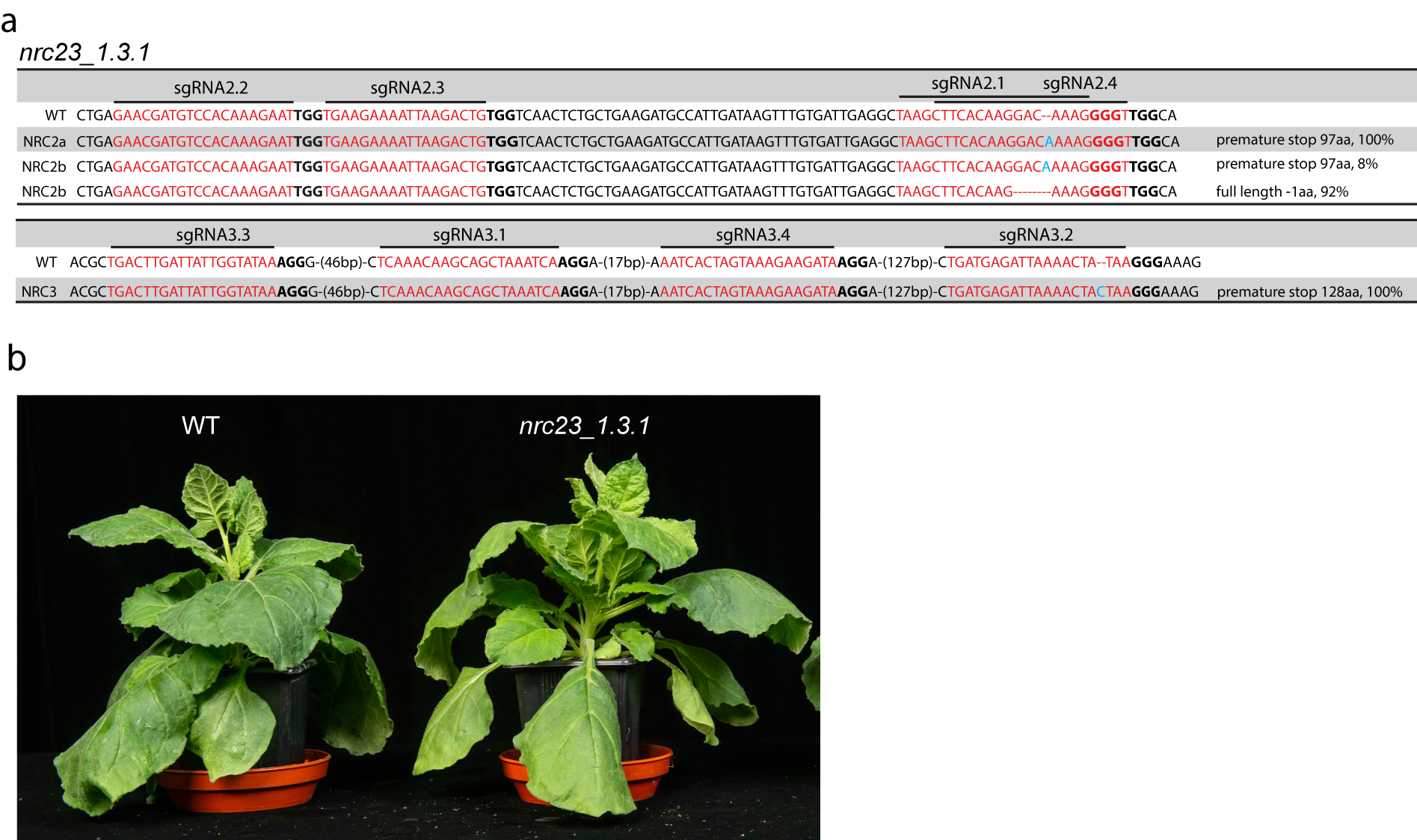

### Fig. S4

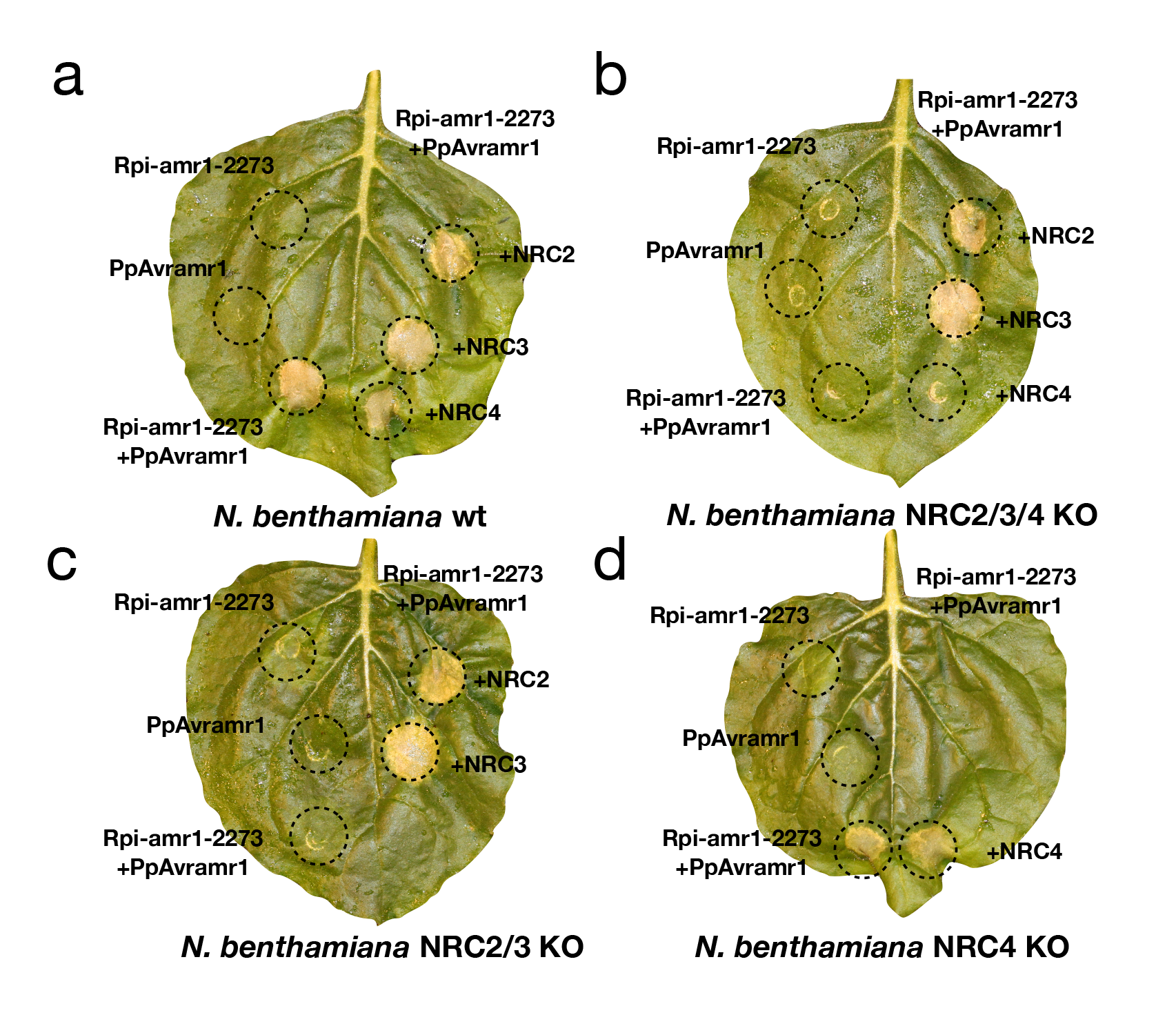

### Fig. S5

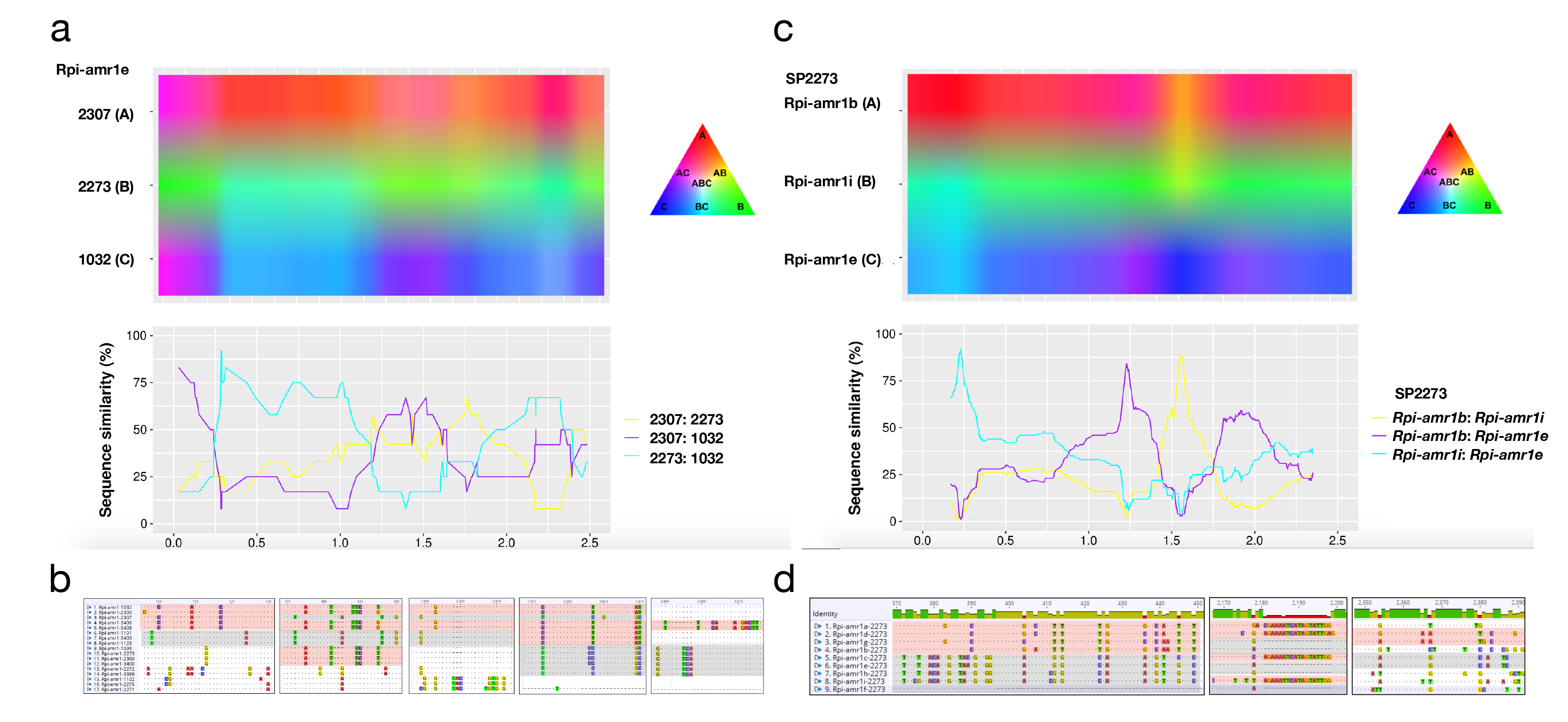

### Fig. S6

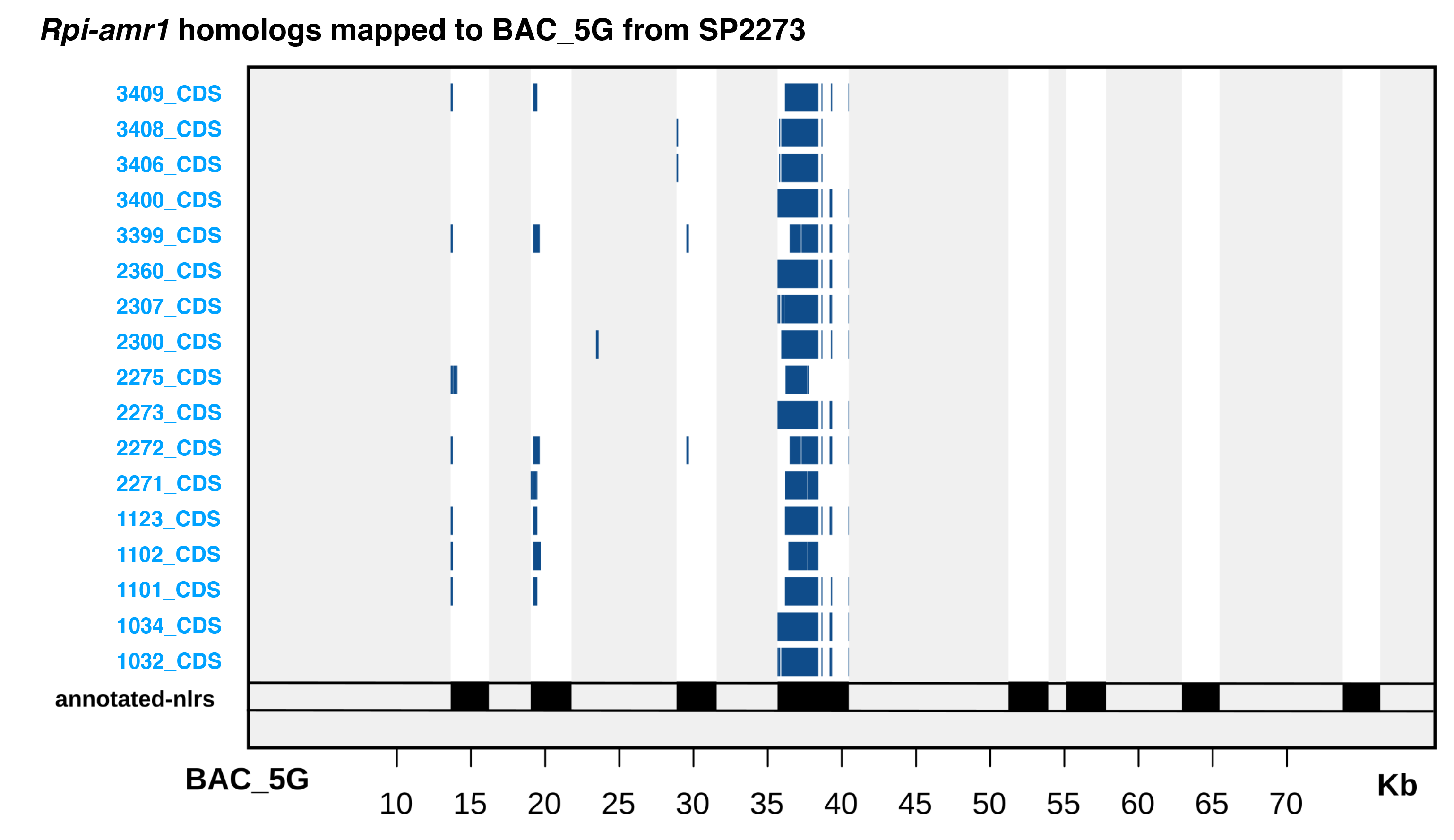
